# Mathematical models disentangle the role of IL-10 feedbacks in human monocytes upon proinflammatory activation

**DOI:** 10.1101/2023.03.24.533939

**Authors:** Niloofar Nikaein, Kedeye Tuerxun, Gunnar Cedersund, Daniel Eklund, Robert Kruse, Eva Särndahl, Eewa Nånberg, Antje Thonig, Dirk Repsilber, Alexander Persson, Elin Nyman, the X-HiDE Consortium

## Abstract

Inflammation is one of the vital mechanisms through which the immune system responds to harmful stimuli. During inflammation, pro and anti-inflammatory cytokines interplay to orchestrate fine-tuned, dynamic immune responses. The cytokine interplay governs switches in the inflammatory response and dictates the propagation of inflammation. Molecular pathways underlying the interplay are complex, and time-resolved monitoring of mediators and cytokines is necessary as a basis to study them in detail. Our understanding can be advanced by *in silico* models which enable to analyze the system of interactions and their dynamical interplay in detail. We, therefore, used a mathematical modeling approach to study the interplay between prominent pro and anti-inflammatory cytokines with a focus on Tumor Necrosis Factor (TNF) and Interleukin 10 (IL-10) in lipopolysaccharide (LPS)-primed primary human monocytes. Relevant time-resolved data were generated by experimentally adding or blocking IL-10 at different time points. The model was successfully trained and could predict independent validation data and was further used to perform *in silico* experiments to disentangle the role of IL-10 feedbacks in acute inflammation. We used the insight to obtain a reduced predictive model including only the necessary IL-10-mediated feedbacks. Finally, the validated reduced model was used to predict early IL-10 – TNF switches in the inflammatory response. Overall, we gained detailed insights into fine-tuning of inflammatory responses in human monocytes and present a model for further use in studying the complex and dynamic process of cytokine-regulated acute inflammation.

## Introduction

Inflammation is central to many common diseases ranging from chronic low-grade inflammatory conditions to diseases of acute, often over-whelming inflammation. The inflammatory process is a sequence of complex and often simultaneous events, including molecular, immunological and physiological mechanisms (1). One of the earliest outcomes of the inflammatory response is the cellular release of soluble mediators, such as proinflammatory cytokines, of which TNF and IL-1β are among the earliest and best described. These are central, pleotropic proinflammatory regulators exerting both autocrine and paracrine signaling, and the primary sources are the myeloid inflammatory cells, including monocytes and certain subsets of macrophages (2,3).

Cellular production and release of these cytokines is transcriptionally dependent on NF-κB activation and translocation of the p65/p50 heterodimer to the nucleus (4). This activation is initiated upon degradation of its inhibitor (IKK) mediated by a number of receptors, including TLR4 (4,5). In addition to transcriptional regulation, TNF is also regulated via stabilization/destabilization of the mRNA pool, while IL-1β is post-translationally activated by the inflammasome complex through enzymatic cleavage (6). Among the most well-known negative regulators of TNF is IL-10, which upon receptor ligation reduces the translocation of p65/p50 heterodimers and the stability of the TNF mRNA pool thus, reducing TNF expression (3). Regarding IL-1β, negative regulation commonly occurs through the soluble receptor antagonist, IL-1Ra, thereby inhibiting the cellular receptor response to IL-1β (7). Hence the balance between proinflammatory cytokines, such as TNF and IL-1β, and anti-inflammatory IL-10 is a critical factor for the outcome of the induced subsequent inflammatory propagation.

Although several studies highlight quantification of proinflammatory cytokines in inflammatory diseases (8), the scarce time points with regards to the inflammatory progression do not allow for a time resolved understanding of e.g., the amplitude and duration dynamics. Studying the balance between the pro and anti-inflammatory cytokines to gain insight into the internal regulation in an episode of inflammation requires time-resolved monitoring of several mediators. Complementary approaches such as *in silico* models help to develop our understanding of the intricate dynamical regulations, where both amplitude of signals and their duration dynamics need to be taken into account to define the regulatory mechanisms (9–11). Such models are built based on the available mechanistic knowledge, i.e., how molecules couple and engage in reactions, and are identified and tuned based on limited sets of time-resolved data. Previous related *in silico* research range from descriptions of detailed single inflammatory pathways (12) to comprehensive maps incorporating multiple layers of inflammatory networks (13). Most of the previous related *in silico* models, based on their questions in hand, describe regulation of inflammatory pathways following either lipopolysaccharide (LPS)-priming (14), TNF stimulation (15–18) or their combinations with anti-inflammatory drugs (19). There have also been several efforts to model these pathways in response to IL-1 (20,21), *S. aureus* (22), and lymphotoxin β receptor (LTβR) signaling (23). However, none of these models have been used to study the role of IL-10 feedbacks in the acute inflammatory phase.

Here, we have developed a mechanistic *in silico* model of the TNF - IL-10 axis, to enable prediction of early switches in the inflammatory response, dictating the propagation of inflammation. To this end, we experimentally augmented or inhibited IL-10 feedback loops at different time points in LPS-primed, primary human monocytes and measured the concentration of TNF, IL-10, IL1-Ra, and IL-1β in a time resolved sample collection routine. This experimental design allowed for thorough investigations of time and amplitude aspects of the interplay using mechanistic modeling. Following data collection, we first created an *in silico* model based on a reaction map, including known biological pathways and interplay between TNF, IL-10, IL-1Ra, and IL-1β, and fitted it to the training experimental data. Second, the data from the IL-10 augmented experimental design was used to validate the model structure. Third, the model was analyzed to find the most important IL-10-mediated feedbacks and the obtained insights were used to introduce a reduced predictive model for the balanced regulation of an LPS-induced inflammatory response, in human monocytes. Finally, we used the validated reduced model to predict the early switches mediated by the IL-10 - TNF interplay in LPS- and IL-10-induced inflammatory response in human monocytes.

## Results

### Data collection: In primary human monocytes, exogenous IL-10 has a time-dependent, dampening effect on the TNF response

Relevant experimental data is essential both for the training of *in silico* models and for model validation, within the modeling cycle (Fig. 1). Here, the experimental data was generated by LPS-priming of a primary human monocyte cell culture to initiate an inflammatory response with focus on the interplay between the proinflammatory cytokine TNF and the anti-inflammatory cytokine IL-10. Four experimental scenarios were designed to capture the interplay in different settings (see **Experimental procedures** - **Data collection** for details). The cells were stimulated with: I) 10 ng/mL of LPS at time point 0h (Control), II) 10 ng/mL of LPS and 10 µg/mL of anti-hIL-10R (a blocking IL-10 receptor antibody) at 0h (IL-10-Block), III) 10 ng/mL of LPS at time point 0h and 100 ng/mL of IL-10, 4h after the administration of LPS (IL-10-4h), and IV) 10 ng/mL of LPS and 100 ng/mL of IL-10, at time point 0h (IL-10-0h). Scenarios I-III were used for model training (Fig. 2B-E) and scenario IV was used for model validation (Fig. 3C). In each scenario, secreted concentrations of the selected cytokines (TNF, IL-10, IL-1Ra, and IL-1β) were measured in cell supernatants at multiple timepoints (0h, 4h, 10h, 24h, 48h). In scenarios III and IV where exogenous IL-10 was added to the cell culture, no quantification of IL-10 was obtained.

**Fig. 1.**
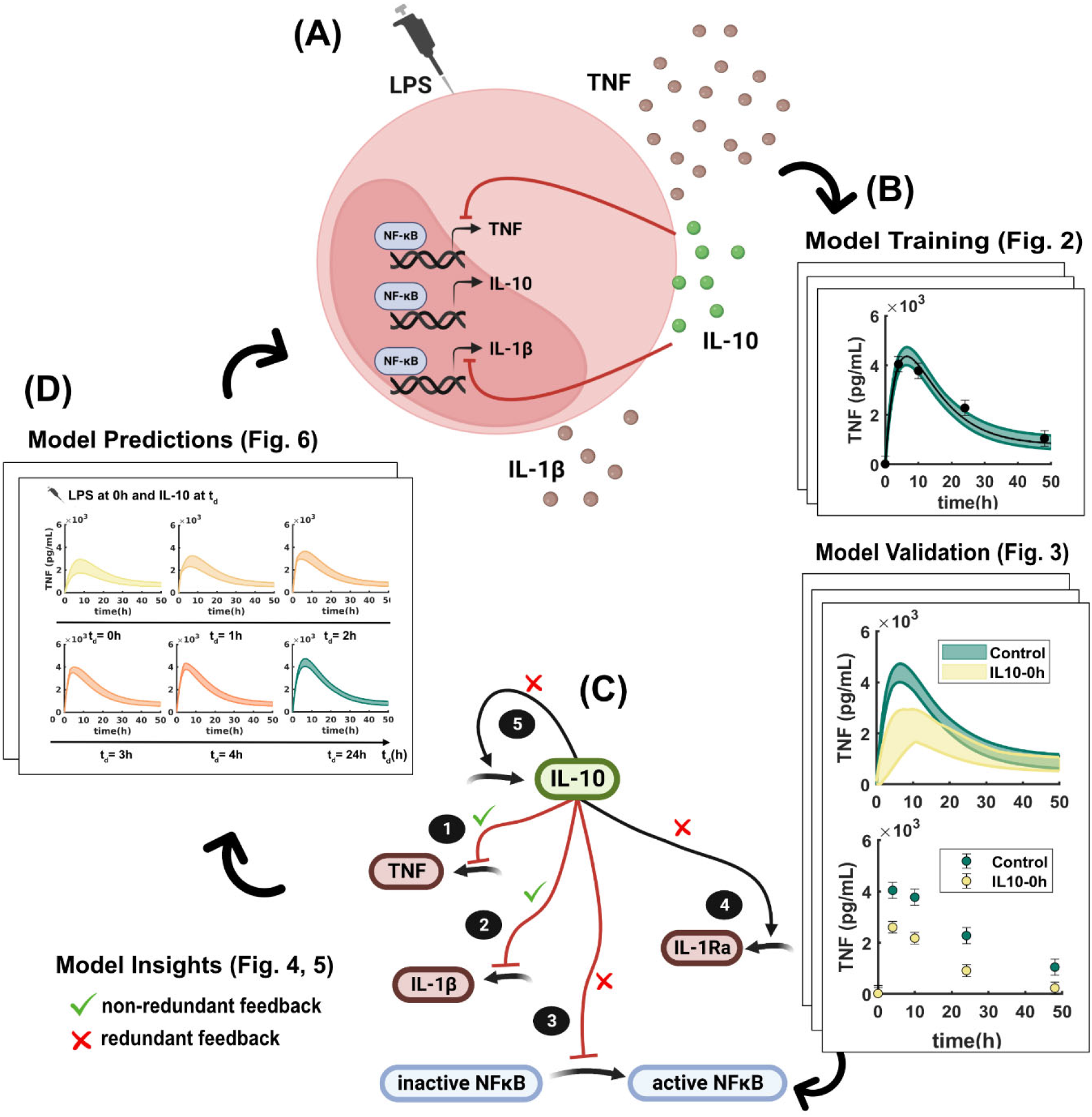
Overview of modeling cycle. (A) A simplified illustration of NF-κB pathway during LPS stimulation of primary human monocytes. LPS stimulates activation of NF-κB, which in turn, regulates expression of pro and anti-inflammatory cytokines. The dynamic interplay between the expressed proinflammatory cytokines and anti-inflammatory cytokine IL-10 is of our interest. Created with BioRender.com (B) *in silico* model development. Biologically relevant experimental data was generated and used to train and validate an *in silico* model of LPS-stimulated primary human monocytes. (C) Model insights. IL-10-mediated inhibition of TNF and IL-1β were found to be necessary for the model to explain the experimental data. Created with BioRender.com (D) Model predictions. The validated *in silico* model predicted a critical 2-hour time window for IL-10 administration post LPS, to damp TNF at the onset of inflammation.

**Fig. 2.**
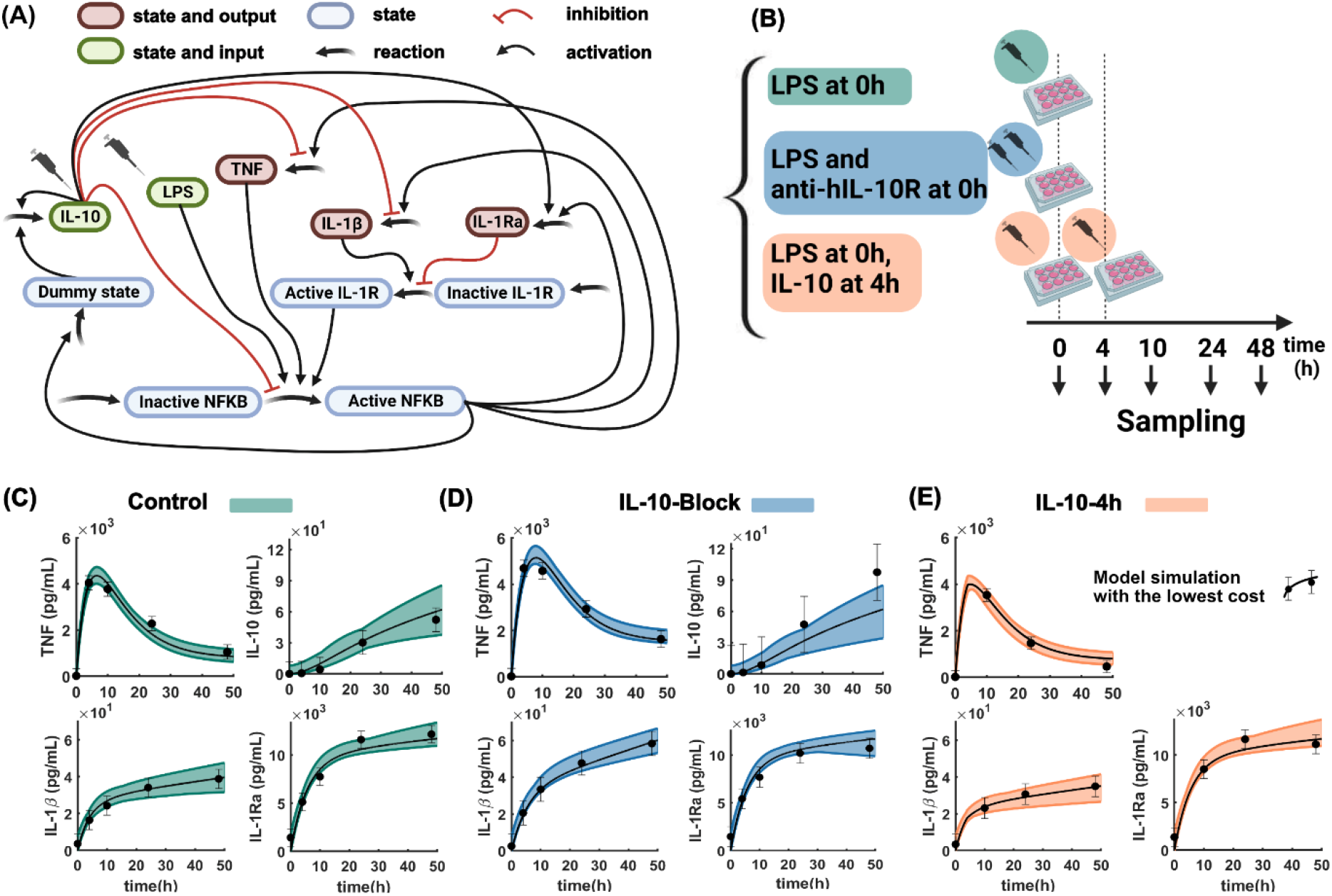
Overview of the training process. (A) Model reaction map. An interdisciplinary team of biomedical experts and mathematical modelers screened the literature to collect available mechanistic knowledge on LPS stimulated monocytes. A reaction map was then proposed based on the collected evidence. Created with BioRender.com (B) Experimental protocols designed to generate data for model training. The primary human monocyte cells were stimulated with LPS at 0h (“Control”), LPS and anti-hIL-10R at 0h (“IL-10-Block”) and LPS at 0h and IL-10 at 4h (“IL-10-4h”). In each scenario, experimental data was collected for four cytokines TNF, IL-10, IL-1β and IL-1Ra, at the specified time points. In the “IL-10-4h” scenario, where exogenous IL-10 was added to the cell culture, no reliable measurement of IL-10 was obtained. Created with BioRender.com (C-E) The *in silico* model was trained based on the collected experimental data. Black dots are the experimentally measured data and error bars are the maximum standard error of the mean (SEM) for each of the measured cytokines. The black solid lines are model simulations with the best agreement to the experimental data. The green, blue and orange areas represent the uncertainty of model predictions for the “Control”, “IL-10-Block” and “IL-10-4h”.

**Fig. 3.**
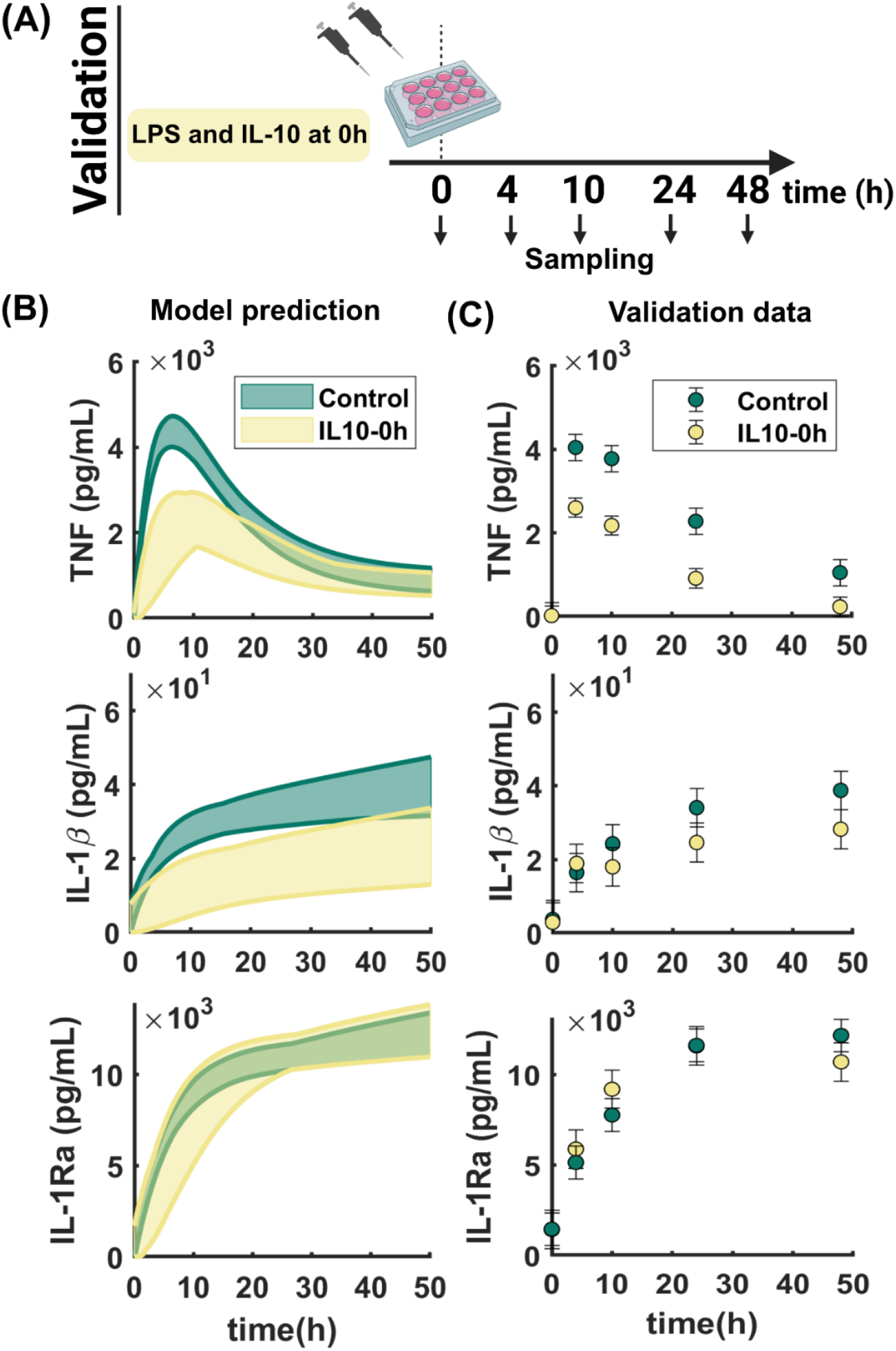
Overview of model validation. (A) The experimental protocol used to generate data for model validation. The primary human monocyte cell culture was stimulated with LPS and IL-10 at 0h (“IL-10-0h”). Experimental data was collected for three cytokines TNF, IL-1β and IL-1Ra, at the specified time points. Due to the addition of exogenous IL-10, no reliable quantification of IL-10 was obtained. Created with BioRender.com (B) Yellow areas are the uncertainty of model predictions for TNF, IL-1β and, IL-1Ra in the “IL-10-0h” scenario, whereas green areas are the model uncertainty for the “Control” scenario. (C) Yellow and green dots are the experimentally measured data, and the error bars are the maximum standard error of the mean (SEM) for each of the measured cytokines in “IL-10-0h” and “Control” scenarios, respectively. The predicted yellow areas overlap the validation experimental data (yellow), indicating successful predictions for all cytokines.

In response to LPS, cells responded with increasing amounts of IL-10, IL-1β, and IL1-Ra over 48 hours, while TNF increased for the first 10-12 hours, whereafter it descended to finally settle at a steady state level (Fig. 2C). Furthermore, IL-10 was secreted with an approximate time delay of 10 hours compared to TNF, IL-1β and IL1-Ra (Fig. 2C, IL-10). Adding exogenous IL-10 together with or 4h later than the LPS-stimulation damped the TNF response compared to that of the control experiment (compare TNF in Fig. 2C, Fig. 2E, and Fig. 3C). In the IL-10-4h scenario, the damping was not as large (7% cf. control, 6h after IL-10 administration, Fig. 2E) as in the IL-10-0h scenario (35% cf. control, 4h after IL-10 administration, Fig. 3C).

Adding exogenous IL-10, dampened IL-1β in both “IL-10-0h” (28.2 ± 11.7 pg/mL, 48h post LPS) and “IL-10-4h” (34.9 ± 13.8 pg/mL, 48h post LPS), compared to the “Control” scenario (38.7 ± 11.6 pg/mL, 48h post LPS) (Fig. 2C, 2E, and Fig. 3C). In addition, endogenous IL-10 exerted inhibition since administration of anti-hIL-10R resulted in an accumulated increase in IL-1β (58.4 ± 16.0 pg/mL, 48 hours post LPS and anti-hIL-10R) compared to the “Control” (Fig. 2D). In conclusion, our experimental data verify the inhibitory role for IL-10 on both TNF and IL-1β of which, the inhibitory effect on TNF appears to be time dependent. To be able to analyze this data further, an Ordinary Differential Equation (ODE)-based model of the TNF - IL-10 axis was built.

### A newly developed mathematical model for the TNF – IL-10 axis can explain training data

We developed an ODE model of the TNF - IL-10 axis, including feedback loops driven by IL-10, TNF, IL-1β, and IL-1Ra. The model was built based on a simplified reaction map of LPS-primed NF-κB signaling pathway (Fig. 2A) (see the **Experimental procedures**). We further successfully trained the model against the generated datasets for scenarios I-III: “Control”, “IL-10-Block” and “IL-10-4h” datasets (Fig. 2C-E). Solid black lines in each subfigure of Fig. 2C-E, represent simulated signals with the best agreement to the training data set, according to the *χ*^2^-test (cost 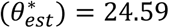 < *χ*^2^(0.05.52) = 69.83, the optimal parameter set 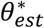 values are presented in Table I). Moreover, the green, blue and orange areas represent the uncertainty of model predictions for the “Control”, “IL-10-Block”, and “IL-10-4h”, respectively (see **Experimental procedures** for details). In short, model simulations for each cytokine are expected to lie in the colored area, indicating agreement to the training data set according to a statistical test (Eq. 13, **Experimental procedures**). This trained model needs to be validated against new data, independent of the training data sets, to prove reliable in terms of prediction.

### Validation: The model successfully predicts independent data collected upon adding exogenous IL-10 at 0h

The “IL-10-0h” data set, in which both LPS and IL-10 were added to the cell culture at 0h (Fig. 3A), was not used in model training and could, therefore, be used to validate the model. The model was stimulated with the same LPS and IL-10 concentrations as in the “IL-10-0h” scenario and model predictions for TNF, IL-1β, and IL-1Ra were maximized and minimized at the measured time points while requiring the agreement to the training data to stay below the threshold for a statistical test (**Experimental procedures**). The obtained model predictions for the “IL-10-0h” scenario were well determined (yellow Fig. 3B) and differed from the “Control” scenario (green Fig. 3B), therefore, useful for model validation. The corresponding experimental data for the “IL-10-0h” scenario are depicted in yellow, and for the “Control” scenario in green (Fig. 3C). Note that the “Control” scenario was used in model training, therefore, only the “IL-10-0h” scenario can be used for model validation. The yellow prediction range was in agreement with the corresponding experimental data for all cytokines indicating that the model successfully captured the damping effect of exogenous IL-10 on both TNF and IL-1β (yellow and green signals in Fig. 3B vs. Fig. 3C). The model also correctly predicted the IL-10 inhibitory effect to be stronger on TNF in the “IL-10-0h” scenario, compared to that of the “IL-10-4h” (compare TNF in Fig. 3B, C to Fig. 2E). Moreover, the model predicted overlaying green and yellow uncertainty ranges for IL-1Ra in the “Control” and “IL-10-4h” scenarios (IL-1Ra in Fig. 3B). The prediction corresponded to the overlapping experimental measurements in these two cases (IL-1Ra in Fig. 3C). The model, now successfully validated, was used to further study the system.

### Investigating the role of IL-10 in the TNF - IL-10 axis and proposing a reduced model including only the necessary IL-10 feedbacks

With the validated model at hand, we investigated the role of IL-10 feedback loops in the TNF - IL-10 axis by eliminating one at a time (Fig. 4, 5). Model-based predictions showed the importance of each of the eliminated feedbacks (Fig. 4A, feedbacks 1-5). The first excluded feedback was IL-10 inhibition of TNF transcription (1 in Fig. 4A). By eliminating this feedback, the model was unable to explain TNF dynamics in the “Control”, “IL-10-0h”, and “IL-10-4h” scenarios (Fig. 4B). Instead, TNF dynamics was predicted to follow that of the “IL-10-Block” scenario in all cases. The second excluded feedback was IL-10 inhibition of IL-1β transcription (2 in Fig. 4A). By eliminating this effect, the model again failed to explain IL-1β dynamics in the “Control”, “IL-10-0h”, and “IL-10-4h” scenarios (Fig. 4C), and instead predicted them to follow the same pattern as in the “IL-10-Block” scenario.

**Fig. 4.**
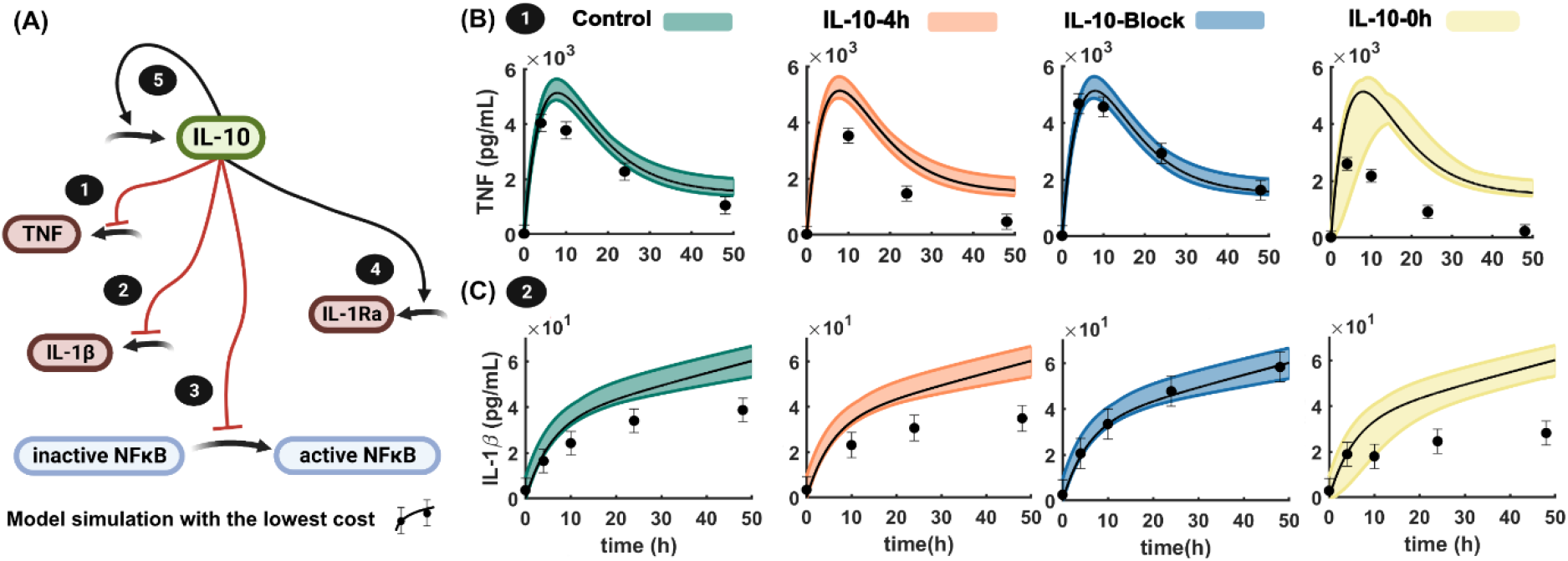
Model analyses. (A) A simplified map illustrating IL-10-mediated regulation of inflammatory cytokines. IL-10 negatively regulates (1) TNF transcription, (2) IL-1β transcription and maturation and (3) NF-κB activation. Moreover, IL-10 positively regulates (4) IL-1Ra and (5) IL-10 expression. Created with BioRender.com (B) Feedback (1) was excluded from the model by setting its corresponding rate parameter to zero. Black dots with error bars are the experimental data and solid black lines and colored areas are the best fit and uncertainty of simulations. The *in silico* model fails to explain TNF experimental data in the absence of IL-10-mediated inhibition of TNF transcription. (C) Feedback (2) was excluded from the model by setting its corresponding rate parameter to zero. The *in silico* model fails to explain IL-1β experimental data in the absence of IL-10-mediated inhibition of IL-1β transcription and maturation.

**Fig. 5.**
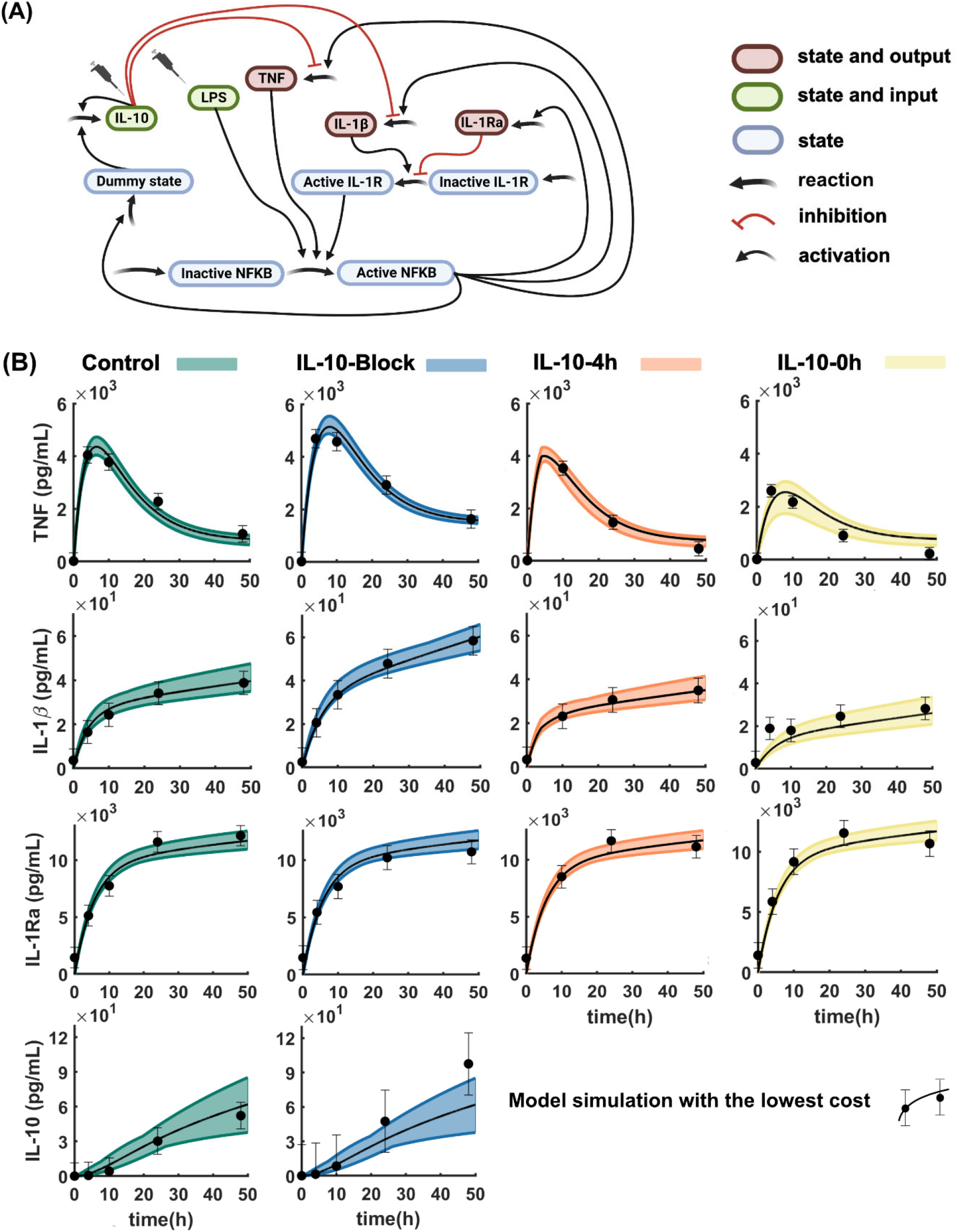
Proposing a reduced model including only the necessary IL-10-mediated feedbacks. (A) a simplified reaction map of the model was proposed by excluding IL-10-mediated negative regulation of NF-κB activation, and positive regulation of IL-10 and IL-1Ra. Created with BioRender.com (B) Feedbacks (3-5) were excluded from the model by setting their corresponding rate parameters to zero. Black dots with error bars are the experimental data and solid black lines and colored areas are the best fit and uncertainty of simulations. The reduced model successfully explains the experimental data in all scenarios.

Next, we proceeded to exclude IL-10 inhibition of NF-κB activation (3 in Fig. 4A), IL-10 positive regulation of IL-1Ra expression (4 in Fig. 4A), and IL-10 positive regulation of IL-10 (5 in Fig. 4A). Unlike negative feedbacks 1 and 2, the model was still capable of explaining the experimental data in all four scenarios in the absence of each feedback 3-5 (not shown). The insights above were used to propose a simplified version of the model that was in agreement with all data and included only the necessary feedbacks (Fig. 5A). The reduced model successfully captured the dynamics observed in all of the measured cytokines (Fig. 5B).

### The model predicts a critical 2-hour time window for IL-10 administration post LPS

By comparing the experimental data collected from the “IL-10-0h” and “IL-10-4h” scenarios, we observed that the inhibitory effect of exogenous IL-10 on TNF is negatively correlated with the time of IL-10 administration (**Results - Data collection**). The reduced model was employed to study the effect of IL-10 administration time on the propagation of inflammation, by monitoring TNF as a potent marker. To this end, 6 different simulations were performed in which the model was stimulated with 10 ng/mL of LPS at 0h, and 100 ng/mL of IL-10 at 0h, 1h, 2h, 3h, 4h, and 24h post LPS (Fig. 6). We observed that the later IL-10 was administered, the less inhibitory effect it enforced on TNF. Up to 2h post LPS-priming, IL-10 substantially impeded TNF, whereas after this critical time span, the simulations roughly followed the pattern of the “Control” experimental data.

**Fig. 6.**
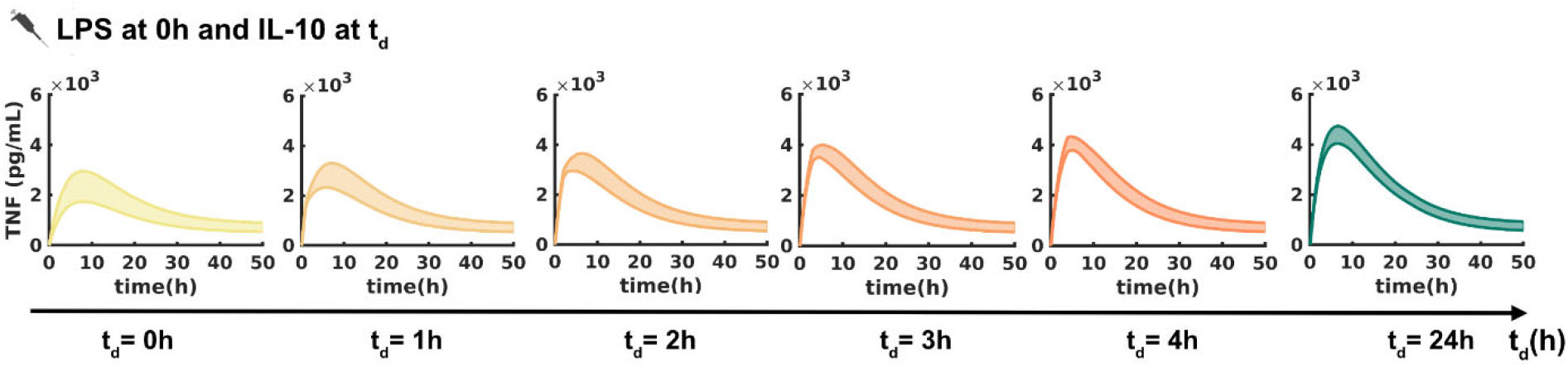
Model simulations to predict the critical time span for IL-10 administration to damp TNF at the onset of inflammation. The model was stimulated with 10 ng/mL of LPS at 0h, and 100 ng/mL of IL-10 at either of t_d_ = 0h, 1h, 2h, 3h, 4h, 24h, post LPS. Each case is depicted in a separate subfigure in which, the uncertainty of prediction is plotted. A critical 2-hour time span for IL-10 administration post LPS is detected through simulations.

## Discussion

By building an *in silico* model, this study investigated the role of IL-10-mediated regulations in LPS-stimulated primary human monocytes. The model was built with a focus on the interplay between pro and anti-inflammatory cytokines and was trained based on data sets collected from biologically relevant experiments in different timing and triggering settings. The model successfully explained the collected data (Fig. 2C-E) and predicted new independent data sets (Fig. 3B). With the validated model at hand, we identified IL-10-mediated negative regulations of TNF and IL-1β as non-redundant, necessary components of balanced regulation of inflammatory responses, in LPS-primed primary human monocytes (Fig. 4, 5). Based on the obtained insights, we reduced the model to include only the necessary IL-10-mediated feedbacks. The reduced model was then used to predict the early IL-10-mediated switches in the inflammatory response, which dictates the propagation of inflammation (Fig. 6).

IL-10 as a potent regulator of inflammation, partly exerts its anti-inflammatory effect through blockage of IKK activation (24), selective formation and translocation of p50/p50, and inhibition of p50/p65 DNA binding (25), via STAT3-SOCS3 axis (26). These effects were modeled by IL-10-mediated inhibition of NF-κB activation in Eq. 3-4 and were assessed to be redundant (Fig. 5). However, IL-10 also tightly regulates both TNF and IL-1β through exclusive regulations. In primary human monocytes, IL-10 inhibits TNF through both transcriptional regulation (27) and destabilization of TNF mRNA (28). These inhibitions were modeled by IL-10-mediated inhibition of TNF in Eq.5 and were assessed to be necessary for the model to explain TNF data in all scenarios (Fig. 4). Moreover, IL-10 regulates IL-1β through two different mechanisms. IL-10 either reduces pro-IL-1β or inhibits maturation of pro-IL-1β by targeting NLRP3 inflammasome activation (29). These regulations take effect through decreasing pro-IL-1β transcriptional or translational rates (30), inhibition of NLRP3 expression (31), enhancing NLRP3 proteasomal degradation (31) or suppressing non-canonical activation of the inflammasome (31,32). These exclusive effects were bundled and modeled by IL-10-mediated inhibition of IL-1β in Eq. 6, which was assessed to be necessary for the model to explain IL-1β data in all scenarios (Fig. 4).

IL-10 further inhibits NF-κB pathway through positive self-regulation (33). IL-10 also positively regulates IL-1Ra, which through IL-1β signaling inhibits NF-κB pathway (34). However, when these IL-10-mediated feedback loops were excluded from the model, simulations still maintained a good agreement with the collected data. The obtained insights on the IL-10-mediated feedbacks and their redundancy, were used to propose a reduced model structure, which only included the necessary IL-10-mediated feedbacks (Fig. 5A, 5B).

We further noticed that exogenous IL-10 exerts a time-dependent inhibitory effect on TNF. TNF in the “IL-10-0h” scenario was damped compared to the “Control” scenario, and even more considerably so when compared to the TNF in the “IL-10-4h” scenario (see **Results-Data collection**). The observed variance in the IL-10 inhibitory effect, which was successfully captured by the model, can be justified by the TNF positive regulation of NF-κB activation. The sooner TNF expression is damped, the less TNF is expressed, resulting in weaker TNF-mediated positive regulations of the pathway.

We used the validated, reduced model to further study the correlation between IL-10 inhibitory effect on TNF and the time of IL-10 administration (Fig. 6). We detected a critical 2h time window during which IL-10 administration substantially diminished TNF as a potent marker of inflammation. After this critical time span has passed, despite addition of IL-10, TNF roughly follows the same pattern as in the “Control” experimental data, indicating a state of less sensitivity to exogenous IL-10. Thus, the later the administration of IL-10, the weaker the inhibitory effect on TNF. This predicted behavioral pattern is of no surprise as TNF is mostly expressed in the early hours post LPS-priming, with a peak around 8-10h (see TNF experimental data in Fig. 2C). This observation indicates a limited time window for interference and the necessity of a tightly controlled dynamics for IL-10.

There was a 10-12h delay in IL-10 expression compared to the expression of other cytokines (Fig. 2C). Delayed expression of IL-10 has previously been observed to be associated to the autocrine/paracrine loop of type I interferons (IFN) in LPS-primed macrophages (35,36). This delayed anti-inflammatory response is suggested to guarantee attenuation of inflammation in a well-timed manner (35). While building the model herein, we noticed that without taking this delay into account, the model could not explain the experimental data. A dummy state, which needs to be identified regarding its biological basis, was hence added to the structure to model the delay (see Eq. 8 in Model Formulation in the **Experimental procedures**).

Another consideration in the model structure was the relation between endogenous presence and exogenous addition of IL-10. In both “IL-10-0h” and “IL-10-4h” scenarios, a high dose of exogenous IL-10 (100 ng/mL) was added experimentally to the cell culture to ensure a locally high IL-10 concentration with similar effect as endogenous IL-10. We did not model the local and global concentrations of IL-10 explicitly, instead we included a variable for the IL-10 effect that was dependent on the sum of exogenous and endogenous IL-10. This IL-10 effect was modeled to be saturated with Michaelis Menten kinetics with a Michaelis constant with an estimated best value of 6.6 pg/mL (see Table 1 and Eq. 2 in Model Formulation in the **Experimental procedures**). Furthermore, in the “IL-10-0h” and “IL-10-4h” scenarios, due to the addition of high levels of exogenous IL-10, no reliable experimental quantification of IL-10 could be obtained. This limitation is due to the linear range of detection.

**Table 1:** The optimal parameter set.

| Parameter | Value | Parameter | Value |
| --- | --- | --- | --- |
| $k_{act1}$ | 8.364e+0 | $k_{feed5}$ | 1.309e+0 |
| $k_{act2}$ | 24.727e-3 | $k_{feed6}$ | 4.616e-3 |
| $k_{act3}$ | 3.549e+0 | $k_{sat1}$ | 6.593e+0 |
| $k_{act4}$ | 12.718e-3 | $k_{sat2}$ | 40.824e-3 |
| $k_{act5}$ | 3.437e+0 | $k_{sat3}$ | 3.182e-3 |
| $k_{act6}$ | 2.586e-3 | $k_{deg1}$ | 16.033e-3 |
| $k_{act7}$ | 74.320e-3 | $k_{deg2}$ | 182.866e-3 |
| $k_{act8}$ | 176.327e-3 | $k_{deg3}$ | 108.570e-3 |
| $k_{act9}$ | 1.222e-3 | $k_{deg4}$ | 863.685e-6 |
| $k_{act10}$ | 12.142e-6 | $k_{deg5}$ | 10.559e-3 |
| $k_{feed1}$ | 12.501e-6 | $k_{deg6}$ | 122.690e-3 |
| $k_{feed2}$ | 4.872e-3 | $k_{deg7}$ | 12.344e-3 |
| $k_{feed3}$ | 3.273e+0 | $k_{deg8}$ | 15.999e-3 |
| $k_{feed4}$ | 1.017e+0 | $k_{deg9}$ | 3.141e-3 |

With the reduced model, we showed that not all included interactions were necessary to explain data from the TNF −IL10 axis. More specifically, we could exclude interactions 3-5 in Fig. 1, i.e., a negative regulation of activation from IL-10 on NF-κB activation, and positive regulations on expression from IL10 to itself and to IL-1Ra. Important to consider is that these interactions have only been shown to be redundant when using the data in the present study, and their importance, even for the same system, can still be proven using other sets of data.

In conclusion, we used a mechanistic, *in silico* model to investigate the TNF - IL-10 interplay in primary human monocytes. The framework of a mechanistic model allows for merging qualitative and quantitative knowledge which are manifested in the model structure and parameter values, respectively. The framework is also flexible to adjustments against new knowledge which facilitates gradual expansion of models. Although mechanistic models are relatively minimal in the sense that they only include reactions related to the question in hand, they are exceptionally relevant for the biological applications. In these applications, we are equally interested in “predicting” the outcome as well as in understanding “how and why” a certain outcome occurs. The ability of a mechanistic model to give insight into the underlying mechanisms, rather than performing as a plain black box predictor, makes it a relevant modeling method in the field of cytokine-regulated inflammation.

## Experimental procedures

### Data collection

### Isolation of primary human monocytes

Primary human monocytes were isolated from buffy coats prepared from healthy blood donors. The peripheral blood monocular cells (PBMCs) were separated by density gradient centrifugation using Lymphoprep (Axis-Shield, Oslo, Norway). The CD14^+^ cells were isolated by magnetic sorting using CD14 MACS microbeads (Miltenyi Biotech, Bergisch Gladbach, Germany) according to manufacturer’s protocol. The isolated CD14^+^ cells were cultured in Dulbecco′s Modified Eagle′s Medium (BE12-614F, DMEM, BioWhittaker, Lonza, Basel, Switzerland) supplemented with 10 % human AB+ serum (pooled from five healthy donors), L-glutamine (10 mM), sodium pyruvate (10 mM) and glucose (4.5 g/L) (all from Gibco, Thermo Fisher Scientific, Waltham, MA).

The study was conducted in accordance with the ethical guidelines of declaration of Helsinki, and in agreement with the ethical policy at Örebro University Hospital, Sweden. The blood samples were coded by the Department of Transfusion Medicine at Örebro University Hospital, and the only information revealed to the researchers was gender and year of birth; thereby preventing data to be traced back to a certain individual. The blood was withdrawn at the time for blood donation, putting no extra harm or risk to the donors. The study therefore did not require ethical approval according to paragraph 4 of the Swedish law (2003:460) on Ethical Conduct in Human Research as confirmed by the Swedish Ethical Review Authority (2022-05214-01).

#### Cell culture experiments

Isolated primary monocytes from each donor were seeded (1 × 10^6^ cells/mL) in culture flasks (Sarstedt, Nümbrecht, Germany) in media supplemented with 10 % human AB+ serum and incubated overnight. On the following day, cells were washed twice with warm PBS before being treated with trypsin. Detached cells were then resuspended into 5 × 10^5^ cells/mL in fresh media and reseeded into a 96-well plate with 5×10^4^ cells per well for each time point. Following one hour of resting, cells were stimulated with 10 ng/mL of LPS (LPS-B5 (*E. coli* serotype 055: K59(B5) H-), Invivogen, San Diego, CA) and supernatants were collected at designated time points.

For the experiments using recombinant human IL-10 (rhIL-10) and anti-human IL-10 receptor antibody (anti-hIL-10R), cells were prepared and seeded in 96-well plates using the same procedure as described above. This was followed by addition of 10 ng/mL LPS with 10 µg/mL of anti-hIL-10R (Biolegend, San Diego, CA), whereas 100 ng/mL rh IL-10 (PeproTech, Cranbury, NJ) was added either at the same time as LPS (0h) or 4h after. Cells treated with only LPS, anti-hIL-10R, isotype control for anti-hIL-10R, or LPS with anti-hIL-10R were used as controls. Supernatants were collected at 0h, 4h, 10h, 24h and 48h and stored at −80°C.

#### Quantification of immune mediators

The collected supernatants were then centrifuged to eliminate cells and debris, prior to measurements. Concentrations (LLOQ) of IL-10 (0.196 pg/mL), TNF (0.181 pg/mL), IL-1Ra (0.426 pg/mL), IL-1β (0.258 pg/m) were measured using a customized U-Plex kit (Meso Scale Discovery, Rockville, MD) detected by electrochemiluminescence in Meso QuickPlex SQ 120 (Meso Scale Discovery).

All analyses were performed according to manufacturer’s instructions. Samples with a coefficient of variation (CV) > 20 % were excluded and those detected below the lower limit of quantification were given an arbitrary value corresponding to half of the lowest standard concentration.

### Model formulation

We used the generated experimental data as well as the available knowledge in the literature, to build an *in silico* model of the TNF - IL-10 interplay during inflammation. The model was built based on a system of nonlinear ordinary differential equations in MATLAB R2018b (MathWorks, Natick, MA), and was numerically solved using the IQM toolbox (IntiQuan GmbH, Basel, Switzerland, https://iqmtools.intiquan.com/) (37). Here, we will explain how the equations are formulated based on both the literature as well as our experimental settings. The corresponding interaction map (Fig. 2A) contains most interactions, but we have excluded interactions corresponding to degradation of model states for illustrative purpose. All model states degrade over time. We have also simplified the inputs in the interaction graph for the same reason. The inputs are described in detail below.

In all experiments, we exposed monocytes to LPS, which activates NF-κB transcription factor and regulates expression of our cytokines of interest. We assumed LPS to maintain a constant concentration throughout 48h, when the data was collected (Eq.1):

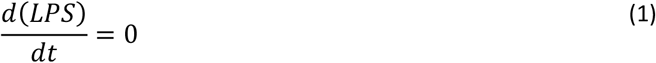

Moreover, in both “IL-10-0h” and “IL-10-4h” scenarios, monocytes are stimulated with exogenous IL-10. IL-10 is expressed endogenously in monocytes as well and therefore, the overall IL-10 effect was modeled by Michaelis Menten kinetics as in Eq. 2:

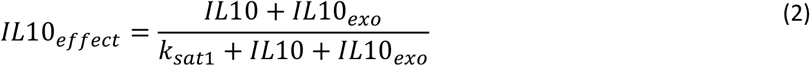

where, *IL*10 and *IL*10_*exo*_ represent endogenous and exogenous IL-10, respectively. A unique Michaelis constant *k*_*sat*1_ was assumed for all IL-10-related regulations and was allowed a free range of [1, 1*e*5] during optimizations.

Besides LPS, TNF and IL-1β positively regulate NF-κB activation, while IL-1R is tightly regulated by the downstream of the NF-κB pathway. To account for the reactions regulating IL-1R and also to avoid complicating the model, we defined IL-1R as a state which encapsulates the effect of IL-1β - IL-1R binding on NF-κB activation. Furthermore, we used Eq.2 to model negative regulation of NF-κB activation by IL-10 (Eq. 3-4):

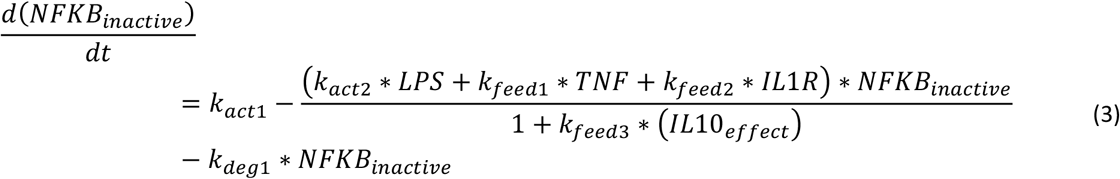

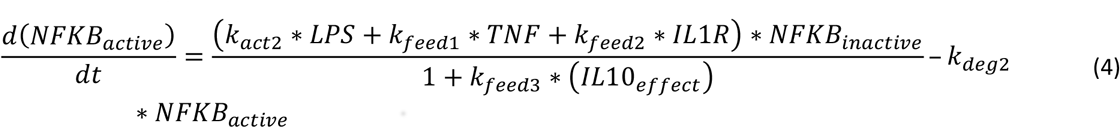

Where, “*NFKB*_*inactive*_”, “*NFKB*_*active*_”, “*TNF*” and “*IL*1*R*” are model states. Moreover, throughout the equations, *k*_*degi*_ are the degradation rate parameters, *k*_*sati*_ are the Michaelis constants and, *k*_*acti*_ and *k*_*feedi*_ are the growth rate parameters for the positive regulations and feedbacks, respectively. “*NFKB*_*inactive*_” is generated by the unknown constant parameter *k*_*act1*_ as a source, and then turns to its active form “*NFKB*_*active*_” through positive regulations mediated by *k*_*actz*_ (*LPS*), *k*_*feed*1_ (*TNF*) and *k*_*feed*2_ (*IL*1*R*), which are formulated based on the mass action law. The activated NF-κB transcription factor would in turn, lead to the expression of TNF, IL-1β and IL-1Ra, while IL-10 negatively regulates TNF and IL-1β and positively regulates IL-1Ra expression (Eq. 5-7):

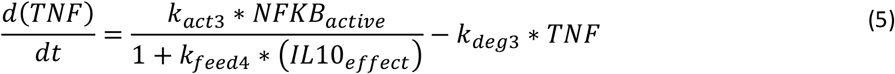

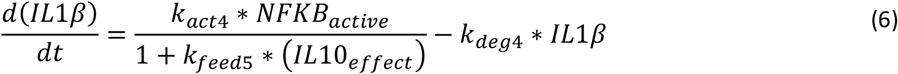

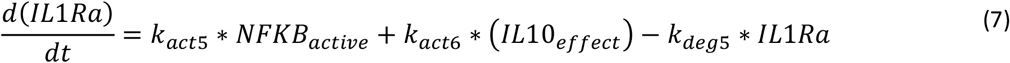

Activated NF-κB also regulates a delayed expression of IL-10, compared to that of TNF, IL-1β and IL-1Ra (see **Data Collection** in **Results**) (Eq. 8-9). To model the observed delay, we introduced a dummy state, “*Dummy*_*state*_”, which could represent any of the intermediate molecules involved in the NF-κB - regulated IL-10 expression, after the activation of NF-κB up until expression of IL-10. The delay was introduced to the system through a Michaelis-Menten kinetics (Eq. 8). Moreover, IL-10 also positively regulates its own expression through the transcription factor Stat3 (33) (Eq. 9).

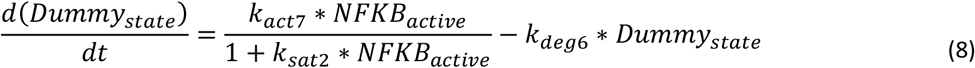

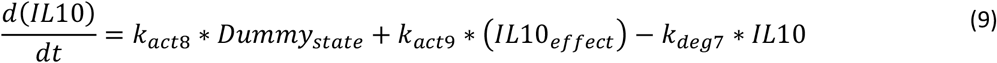

Eventually, we defined two states “*IL*1*R*_*inactive*_” and “*IL*1*R*”, which represent IL-1R inactive and active forms. “*IL*1*R*_*inactive*_” is generated by the parameter *k*_*act10*_ as the source and then binds to *IL*1*β*, while *IL*1*Ra* negatively regulates the binding (Eq. 10–11). Activated IL-1R encapsulates the binding effect of IL-1β to IL-1R on NF-κB activation (see Eq. 3–4).

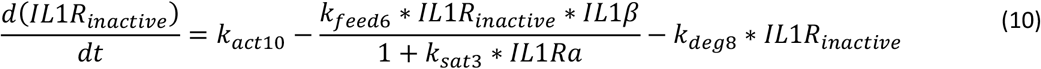

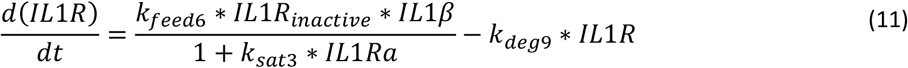

In these equations, *IL*10 and *LPS* are both model states and inputs, while *TNF, IL*1*β, IL*1*Ra* and *IL*10 are both model states and outputs. In the “IL-10-block” scenario, where we needed to block IL-10, we set the parameters *k*_*feed*3_, *k*_*feed*4_, *k*_*feed*5_, *k*_*act*6_, *k*_*act*9_ to zero, to eliminate the IL-10 effect and model the block.

### Parameter Estimation

To train the model, an unknown parameter vector consists of all the above-mentioned rate, feedback, saturation and degradation constants was created and estimated to minimize a cost function of the form *J*:

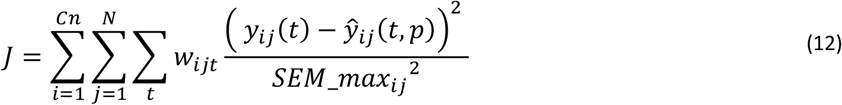

Where, *i=1…Cn* represents different training experimental scenarios, *j=1…N* is the number of measured cytokines in each scenario, and *t* is the time of measurements. Moreover, *y*_*ij*_ (*t*) is the experimentally measured concentration of cytokine *j*, in scenario *i*, at time *t, ŷ*_*ij*_(*t,p*) is the corresponding model simulation as a function of time *t* and parameter vector *p*, and *SEM* - *max*_*ij*_ is the maximum standard error of the mean (SEM) for each of the measured cytokines *j* in scenario *i*. Moreover, *w*_*ijt*_ is the weight parameter used for penalizing the cost of cytokine *j*, in scenario *i*, at time *t*. Model time is assumed to be in hours, and concentrations are in pg/mL, to match units of the measured experimental data.

MATLAB particle swarm (38) and simulannealbnd - simulated annealing algorithm (39) - as well as the simplexIQM - Downhill Simplex Method-from the IQM toolbox were used to search the parameter space, during optimization. Before each optimization iteration or simulation step, the model was run for 9e+06 time units, starting from a zero initial condition to reach its steady state. All parameters were allowed a free range of [1e-5,1e+2] during optimization, unless specified otherwise. To evaluate the models, the *χ*^2^ test (with the significance level of 0.05 and degrees of freedom equal to the number of data points) was applied to assess the agreement between model simulations and the experimental data sets.

### Uncertainty Estimation

The prediction profile likelihood approach (40) was used to determine the uncertainty of predictions. We maximized and minimized model predictions at each measured time point, through solving independent constrained optimization problems. Throughout the optimizations, model simulations were required to maintain their agreement to the training data set by keeping the training cost (Eq. 12) below a threshold of the form (Eq. 13):

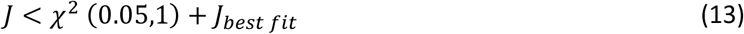

Where, *J*_*best fit*_ is the minimum cost corresponding to the best fit. The constrained optimization problem was solved as implemented in (41), by relaxing the constraint to a penalty term of the form Eq. 14 in case Eq. 13 is not satisfied:

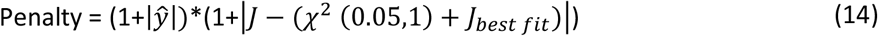

Where, *ŷ* is the prediction value to be optimized.

Model predictions at the measured time points were optimized by running chains of MATLAB hybrid Genetic and fmincon algorithms more than 100 times in total. The collected parameter sets include solutions which maximize or minimize values at each measured time point and for each cytokine. To estimate the uncertainty bounds of predictions between the measured points, the collected parameter sets were used to simulate responses in 48h. The maximum and minimum values of the simulations at each time step (every 6 minutes), were collected and formed the bounds of uncertainty. All simulations were performed using the parallel computing toolbox in MATLAB and a 4 NUMA node processing unit with each node containing an Intel(R) Xeon(R) CPU E5-4627 v3 @ 2.60GHz. A total of 253 GB RAM was accessible during simulations.

## Data availability

DOI will be made available after acceptance.

## Supporting information

No supporting information is available.

### Acknowledgments

We acknowledge scientific support from the Exploring Inflammation in Health and Disease (X-HiDE) Consortium, which is a strategic research profile at Örebro University. Moreover, we want to thank William Lövfors for the helpful discussions on formulation of the optimization problem.

## Author contributions

NN: Methodology, Software, Validation, Formal analysis, Writing - Original Draft, Writing - Review & Editing, Visualization.

KT: Methodology, Investigation, Visualization, Writing - Review & Editing.

GC: Conceptualization, Methodology, Writing - Review & Editing, Supervision, Funding acquisition.

DE: Conceptualization, Methodology, Validation, Resources, Writing - Review & Editing, Supervision, Project administration, Funding acquisition

RK: Conceptualization, Writing - Review & Editing, Supervision, Funding acquisition.

ES: Conceptualization, Writing - Review & Editing, Supervision, Funding acquisition.

EN: Conceptualization, Methodology, Writing - Review & Editing, Supervision, Project administration.

AT: Validation, Investigation, Writing - Review & Editing.

DR: Conceptualization, Methodology, Resources, Writing - Review & Editing, Supervision, Project administration, Funding acquisition.

AP: Conceptualization, Methodology, Validation, Investigation, Resources, Writing - Original Draft, Writing - Review & Editing, Supervision, Funding acquisition.

EN: Conceptualization, Methodology, Validation, Formal analysis, Writing - Original Draft, Writing - Review & Editing, Visualization, Supervision, Project administration, Funding acquisition.

## Funding and additional information

**The X-HiDE Consortium** is funded by the Knowledge Foundation (20200017) and by strategic funding by Örebro University. **GC** acknowledges support from the Swedish Research Council (2018-05418, 2018-03319), CENIIT (15.09), the Swedish Foundation for Strategic Research (ITM17-0245), SciLifeLab National COVID-19 Research Program financed by the Knut and Alice Wallenberg Foundation (2020.0182), the H2020 project PRECISE4Q (777107), the Swedish Fund for Research without Animal Experiments (F2019-0010), ELLIIT (2020-A12), and VINNOVA (VisualSweden, 2020-04711). **EN** acknowledges support from the Swedish Research Council (Dnr 2019-03767), the Heart and Lung Foundation, CENIIT (20.08), åke Wibergs Stiftelse (M19-0449, M21-0030, M22-0027), and the Swedish Fund for Research without Animal Experiments (S2021-0008). The funders had no role in study design, data collection and analysis, decision to publish, or preparation of the manuscript.

